# De novo genome assembly and functional annotation for *Fusarium langsethiae*

**DOI:** 10.1101/2021.04.13.439621

**Authors:** Ya Zuo, Carol Verheecke-Vaessen, Corentin Molitor, Angel Medina, Naresh Magan, Fady Mohareb

## Abstract

**Motivation:** *Fusarium langsethiae* is a T-2 and HT-2 mycotoxins producing *Fusarium* species firstly characterised in 2004. It is commonly isolated from oats in Northern Europe. T-2 and HT-2 mycotoxins exhibit immunological and haemotological effects in animal health mainly through inhibition of protein, RNA and DNA synthesis. The development of a high-quality and comprehensively annotated assembly for this species is therefore essential in providing the molecular understanding and the mechanism of T-2 and HT-2 biosynthesis in *F. langsethiae* to help develop effective control strategies.

**Results:** The *F. langsethiae* assembly was produced using PacBio long reads, which were then assembled independently using Canu, SMARTdenovo and Flye; producing a genome assembly total length of 59Mb and N50 of 3.51Mb. A total of 19,336 coding genes were identified using RNA-Seq informed *ab-initio* gene prediction. Finally, predicting genes were annotated using the basic local alignment search tool (BLAST) against the NCBI non-redundant (NR) genome database and protein hits were annotated using InterProScan. Genes with blast hits were functionally annotated with Gene Ontology.

**Data availability:** Raw sequence reads and assembled genome can be downloaded from: GenBank under the accession JAFFKB000000000

## 1 Introduction

*Fusarium langsethiae* is a fungus belonging to the family *Nectriaceae*. It commonly infects ripening oats without showing any visible symptoms especially in Northern Europe, and contaminates the grains with the type A trichothecenes, T-2 and HT-2 (Torp and Nirenberg, 2004; Kalantari, 2010). These mycotoxins mainly inhibit proteins, RNA and DNA synthesis leading to immunological and haematological effects (CONTAM, 2011).

Originally, this species was considered to be a “powdery *F. poae*”. However, subsequently it was shown to be a separate species and classified as *F. langsethiae* in 2004 by Torp and Nirenberg (Torp and Nirenberg, 2004). More than 300 *Fusarium* species exist (Summerell, 2019) with different ecophysiological responses to parameters such as temperature and water availability. *F. langsethiae* is a relatively slow colonizer of temperate cereals although the ability to produce mycotoxins may provide some competitiveness in the phyllosphere microbiome (Medina and Magan, 2010, 2011; Kokkonen, 2012; Verheecke, 2020). Some other more economically important *Fusarium* species, e.g. *F. graminearum*, have received more attention as they produce visible symptoms in wheat (Fusarium head blight) and have thus been widely sequenced and analyzed previously (King, 2015). However, many of these studies have focused on the toxin biosynthesis process (Rep, 2010; Mehrej, 2011). Some of these studies use sequencing methods to analyse protein structure or toxin-related biosynthetic gene clusters. However, most of the *Fusarium* sequences comes from short reads platforms, which may have thus excluded some important functional gene clusters and their proteins. The aim of this work was to produce a high quality, deep coverage genome assembly for *F. langsethiae* through a long-reads assembly strategy using the PacBio platform. Further analysis including identification of the clusters related to biosynthesis of T-2 and HT-2 mycotoxins was also examined. The relatively small length of fungal genomes makes them ideal candidates for long-reads assembly, since it is possible to achieve deep read coverage, allowing a higher degree of continuity with no noticeable implication on the sequencing costs.

## 2 Methods

### 2.1 Sequencing Data

To obtain long sequence fungal DNA, the protocol from Bacha (2015) was used. Briefly, 3-day-old colonies of *F. langsethiae* Fe2391 grown on Potatoes Dextrose Agar (PDA) were harvested and frozen in liquid nitrogen. The mycelia were incubated for 10 min at 50°C in a modified lysis buffer (1% of hexadecyl-trimethyl-ammonium bromide, 100mM pH8 EDTA, 1.4 M NaCl, 20mM pH8 Tris-HCl). The DNA was then extracted 3 times in phenol:chloroform prior to precipitation in isopropanol. The pellet was resuspended with 25U of RNAse prior to sending for sequencing. The DNA size (~20kb) was validated by gel electrophoresis. The samples were sent to Novogene, China, which generated raw sequencing data using the PacBio^®^ Sequel platform.

### 2.2 Genome Assembly

In order to achieve the best assembly quality possible, three separate assemblers were used to process the raw *F. langsethiae* data, namely Canu v1.8, SMARTdenovo, and Flye v2.4.2. Canu (Koren, 2017) has three phases in its pipeline: correction, trimming and assembly. Since Canu is sensitive to the sequences’ genome size, it requires a parameter called ‘error rate’, which refers to the percentage of difference between the two reads in an alignment. The genome size parameter was set to 37,500,000 for this study, based on the length of the assembly publicly available for this species. According to Canus’ user guide, the parameters of error rate should be adjusted according to the coverage and data type of the raw data. Therefore, the error rates were set to 0.045, 0.055, 0.065, 0.075, 0.080, 0.085 and 0.095; this kept the correction, trimming and assembling stages identical. The second assembler used was SMARTdenovo (available at https://github.com/ruanjue/smartdenovo). This tool directly utilises reads from raw read alignments without correction or trimming phases, and it provides its own polishing methods to generate accurate consensus sequences. For the present study, the raw data were directly processed by the SMARTdenovo.pl script with all parameters set to default. The last, and the best performing assembler of the three was Flye (Kolmogorov, 2019), previously called A-Bruijn. It uses a repeat graph as its core data structure and utilises raw data in FASTA or FASTQ format from PacBio^®^. Flye outputs polished contigs with an error rate less than 30% by default. To run the assembly, the genome size was set to 37,500,000 and other parameters were set to default.

A total of five draft assemblies were generated using all three assemblers. QUAST, a quality assessment tool for genome assemblies (Gurevich, 2013), was used to examine the basic quality among the contig assemblies by comparing their total length, longest contig and N50 number. The best assembly output was identified according to completeness through orthologs comparison versus the *Saccharomyceta* OrthoDB (Kriventseva et al. 2019) data set using the Benchmarking Universal Single-Copy Orthologs (BUSCO) tool (Robert, 2017). BUSCO results were judged based on the number of complete BUSCO genes, and then the best assembly was used in the next stage.

To further improve the assembly, polishing was performed using Pilon (Walker, 2014). Since short reads were absent for this sample, the WGS sequence reads of sample Fl201059 (Lysoe, 2016) were downloaded from the European Nucleotide Archive and aligned to our assembly using BWA-MEM (Li, 2009). This resulted in an alignment file in BAM format, which was then used by Pilon to perform error correction.

### 2.3 Gene prediction and functional annotation

Two gene prediction methods, GeneID (Blanco, 2002) and Augustus (Stanke, 2008), were used with four separate procedures, GeneID *ab-inito*, Augustus *ab-inito*, Augustus with hints and Augustus with training. For *ab-initio*, Augustus predicted genes using *F. graminearum*, and GeneID predicted genes using *F. oxysporum*. Only Augustus had settings that could be used for hints prediction and training. The hints were created from the cDNA file from Fl201059. For training, the BRAKER pipeline (Gremme, 2013) with the protein coding file from Fl201059 was used; this pipeline contains an automatic training and prediction pathway using GenomeThreader and Augustus.

A basic local alignment search tool (BLAST; Boratyn, 2019) was used to find regions of local similarity between sequences. Gene sequences that had been extracted from prediction tools were output in a GFF format and then made into a FASTA file. Using the blastx command, the FASTA file was compared with the NR nucleotide database. The number of threads was set to 50. The output format of this command was set as BLAST archive (ASN.1). Functional annotation was performed using OmicsBox (available at https://www.biobam.com/). The BLAST hits were imported into OmicsBox to perform Gene Ontology mapping and annotation. InterPro protein signatures and domain hits were obtained using InterProScan5. The output was then imported in OmicsBox and merged with the GO annotation and mapping results.

## 3 Results

Since this species had not been previously assembled using long-reads, different assemblers were compared to achieve a higher quality overall assembly. After comparing each assembly’s statistical information and BUSCO results, the best assembly was polished as the final assembly. Assembly quality metrics for all five chosen are shown in Table 1, which outlines some basic statistical data from the different assemblies. The total assembly lengths obtained from Canu and Flye were relatively closer to each other compared to SMARTdenovo, which had the longest genome length and the largest number of contigs, indicating fragmentation. Most of the GC content results in this study were around 48.4%; the SMARTdenovo assembly was slightly lower. The highest N50 value was achieved by Flye, which was nearly five times longer than the best Canu assembly and 22 times longer than the SMARTdenovo assembly.

**Table 1:** Basic statistic information of draft assembly from Canu, SMARTdenovo and Flye.

| Assembly method | Canu |  |  | SMART denovo | Flye |
| --- | --- | --- | --- | --- | --- |
| Corrected error rate | 0.045 | 0.065 | 0.085 | - | - |
| Contigs | 301 | 301 | 280 | 277,164 | 177 |
| Contigs (>=50kb) | 174 | 174 | 168 | 170 | 60 |
| Total contigs length | 62,453k | 62,451k | 62,870k | 3,850,300k | 59,663k |
| Total contigs length (>=50kb) | 59,418k | 59,416k | 59,802k | 9,340k | 59,004k |
| Longest contig | 2,495k | 2,495k | 2,723k | 77k | 11,601k |
| GC content (%) | 48.46 | 48.48 | 48.44 | 48.21 | 48.43 |
| N50 | 512k | 521k | 614k | 16k | 3,513k |

The draft assembly from Flye achieved 98.3% completeness, as assessed by BUSCO, and had fewer fragmented and missing hits compared with Canu (See Table 2). On the other hand, the SMARTdenovo assembly only had 11 complete hits, probably due to the fragmented state of this assembly. Considering the contigs length, N50and BUSCO results, Flye output was considered the best assembly which was then carried forward for error correction.

**Table 2:**
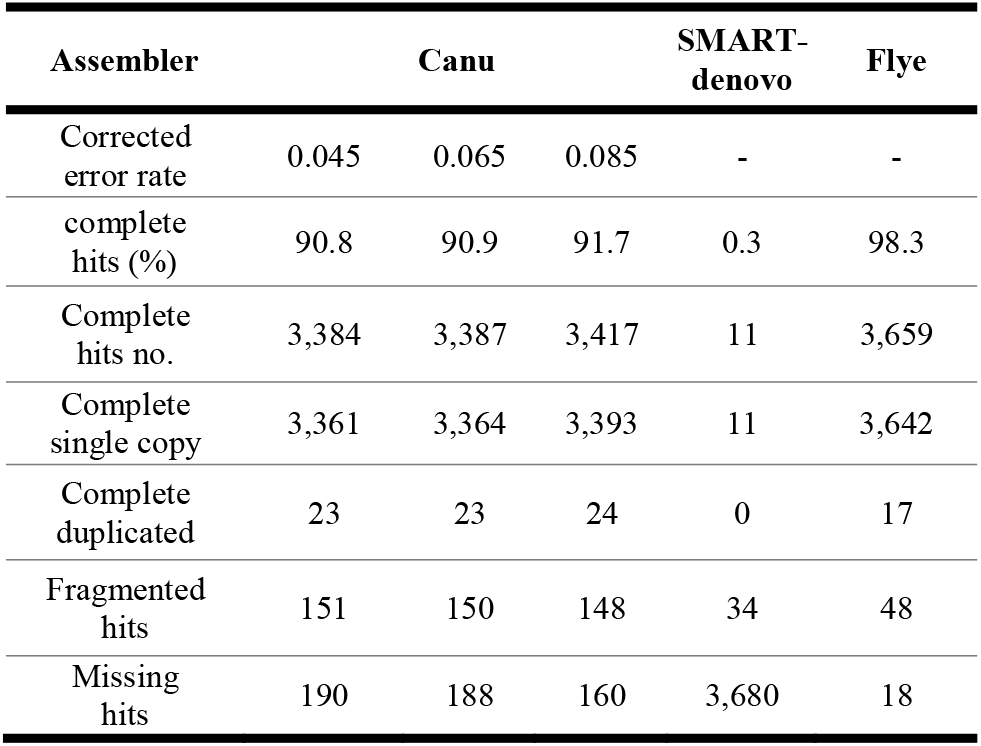
BUSCOs statistic result from Canu, SMARTdenovo and Flye.

In order to improve the assembly quality further, error correction was performed with Pilon based on the previously mentioned publicly available Illumina short reads. Comparing the statistical sequence information and BUSCO results between draft assembly and polished assembly, the length of contigs did not change after hybrid polishing, but the BUSCO results improved from 98.3% to 98.8% (See Table 3). Moreover, 28 of the fragmented genes became either complete (20) or missing (8), which support the fact that Pilon actually removed mis-assemblies from the raw draft. Compared with the previous assembly of another *F. langsethiae* strain (Fl201059), the draft assembly of this study has better contiguity and slightly higher BUSCO results.

To finish the polishing, 23 contigs were detected as mitochondrial hits and were removed from the assembly.

**Table 3:**
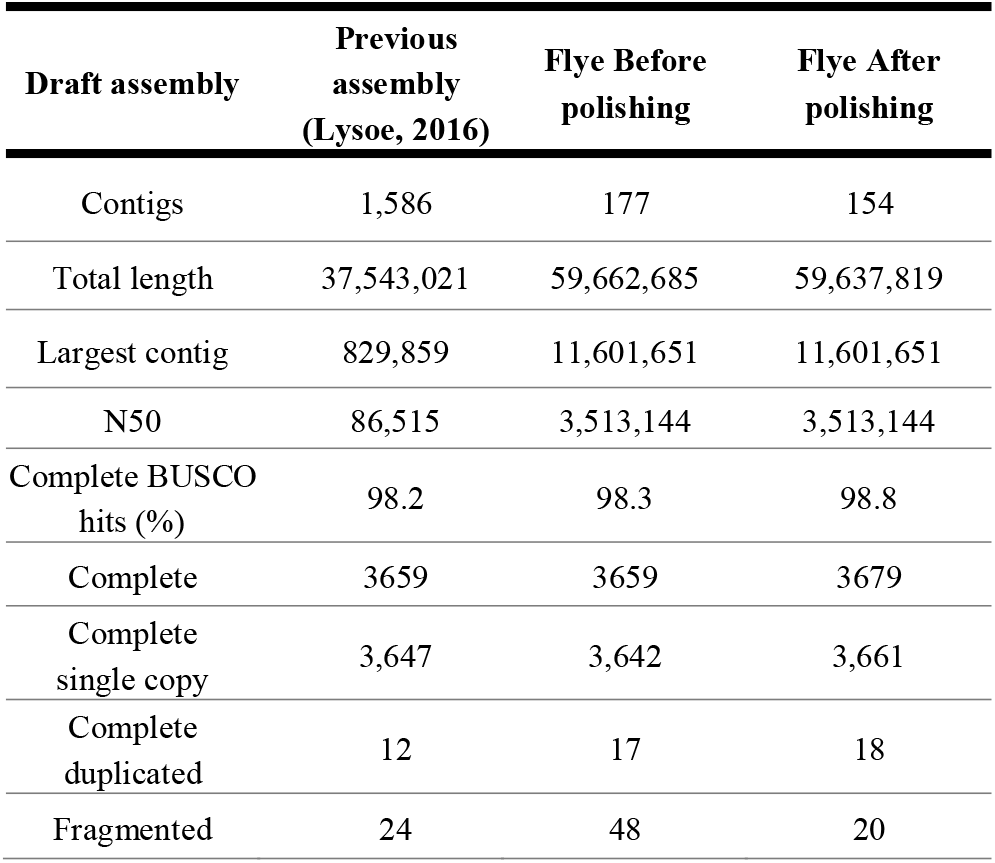
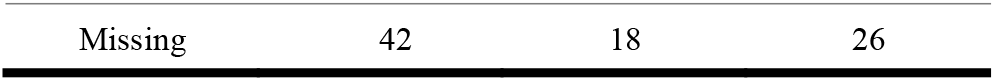
Basic sequence statistic information and BUSCO result comparation between before polishing and after polishing.

| Draft assembly | Previous assembly (Lysoe, 2016) | Flye Before polishing | Flye After polishing |
| --- | --- | --- | --- |
| Contigs | 1,586 | 177 | 154 |
| Total length | 37,543,021 | 59,662,685 | 59,637,819 |
| Largest contig | 829,859 | 11,601,651 | 11,601,651 |
| N50 | 86,515 | 3,513,144 | 3,513,144 |
| Complete BUSCO hits (%) | 98.2 | 98.3 | 98.8 |
| Complete | 3659 | 3659 | 3679 |
| Complete single copy | 3,647 | 3,642 | 3,661 |
| Complete duplicated | 12 | 17 | 18 |
| Fragmented | 24 | 48 | 20 |
| Missing | 42 | 18 | 26 |

A series of gene prediction approaches were followed using different settings or alignment models from related species as shown in Table 4. A hints file was created for Augustus using the cDNA sequence of sample Fl201059 downloaded from EMBL-EBI (Lysoe, 2016). Firstly, the cDNA contigs were aligned to the assembled genome in order to confirm its suitability. A total of 15,280 alignments were obtained, representing 96% mapping results which confirms its suitability to guide the gene prediction process. GeneID in ab-initio mode predicted 21,848 genes, while Augustus predicted a total 16,900 genes. Then, after training, Augustus predicted 19,336 genes.

**Table 4:**
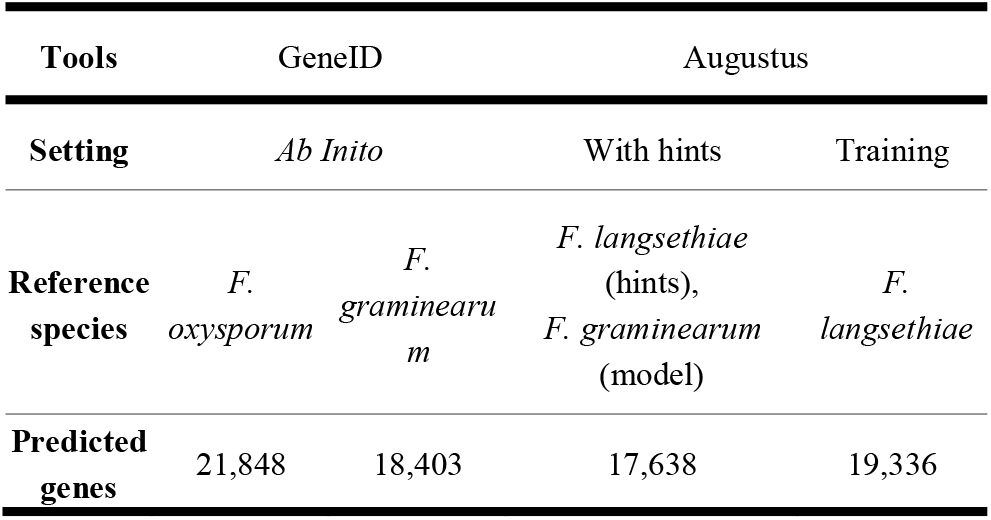
Number of predicted genes from different tools, settings, and reference species.

Following the gene prediction step, a BLAST search was performed of predicted coding genes against the NR database, and the hits were further annotated with GO terms and protein signatures. A total of 19,139 out of 19,336 predicted genes (98.98%) had more than five hits against the NR database. This meant that more than 99% of genes predicted by Augustus were reliable for protein analysis to some extent. Among the hits, most had a similarity percentage higher than 80%; some of them reached 100% similarity with the sequence in the NR database.

More than 50% of the genes in the assembly had top hits compared with *F. langsethiae* itself. Almost all of these were within the *Fusarium* genus. Table 7 shows the top 10 BLAST hits distribution, all of them coming from the *Fusarium* genus. The assembly showed a linkage between *F. langsethiae* and *F. poae*, *F. oxysporum, F. graminearum*, *F. sporotrichioides* therefore indicating a similarity in the metabolic pathways and/or mycotoxin production. As the *F. poae* had the highest number of hits amongst all the *Fusarium* species, it suggests a close relevance between *F. langsethiae* and *F. poae*. Indeed, it should be considered that in papers published before 2004, *F. langsethiae* was considered as “powdery *F. poae”.* It could be inferred that, *F. poae* should thus have the closest linkage to *F. langsethiae*, amongst all the *Fusarium* species. The full chart of mycotoxins and their related proteins is provided in the Supplementary Materials SF1. However, some proteins related to trichothecene and HC-Toxin are shown in Tables 8 & 9 respectively. These tables list the contigs in which each protein was located, as well as the similarity to genes in other species as found via BLAST. The ontology term and ontology ID give a basic description of the protein and its functions.

**Table 7:** Top 20 blast top hits distribution among predicted genes.

| Species | Top hits |
| --- | --- |
| *Fusarium poae* | 50,396 |
| *Fusarium oxysporum* | 33,516 |
| *Fusarium oxysporum f. sp. cepae* | 25,207 |
| *Fusarium graminearum* | 24,162 |
| *Fusarium langsethiae* | 15,412 |
| *Fusarium oxysporum f. sp. cubense* | 14,995 |
| *Fusarium fujikuroi* | 11,795 |
| *Fusarium sporotrichioides* | 11,391 |
| *Fusarium oxysporum f. sp. narcissi* | 11,259 |
| *Fusarium venenatum* | 11,110 |

**Table 8:**
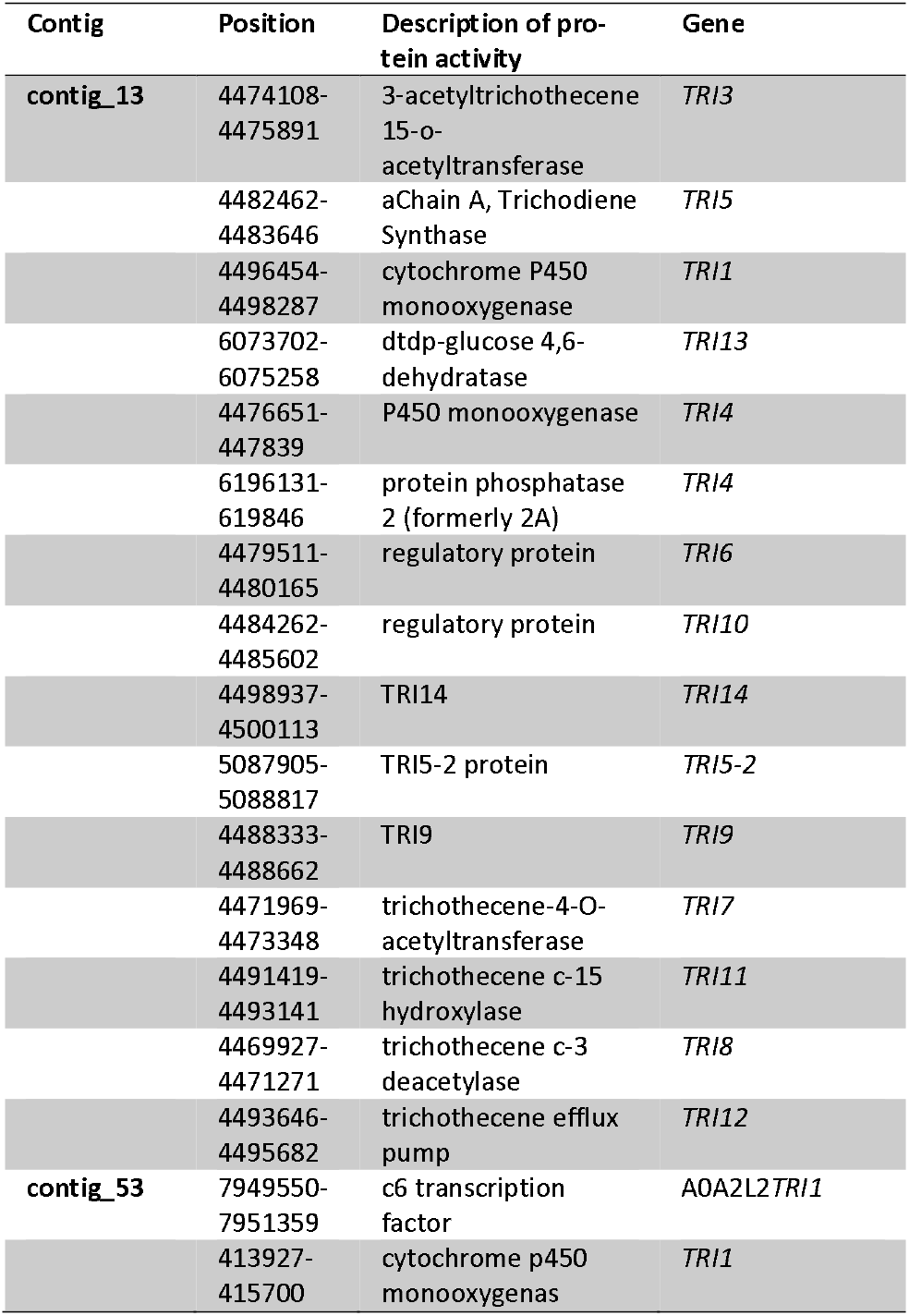
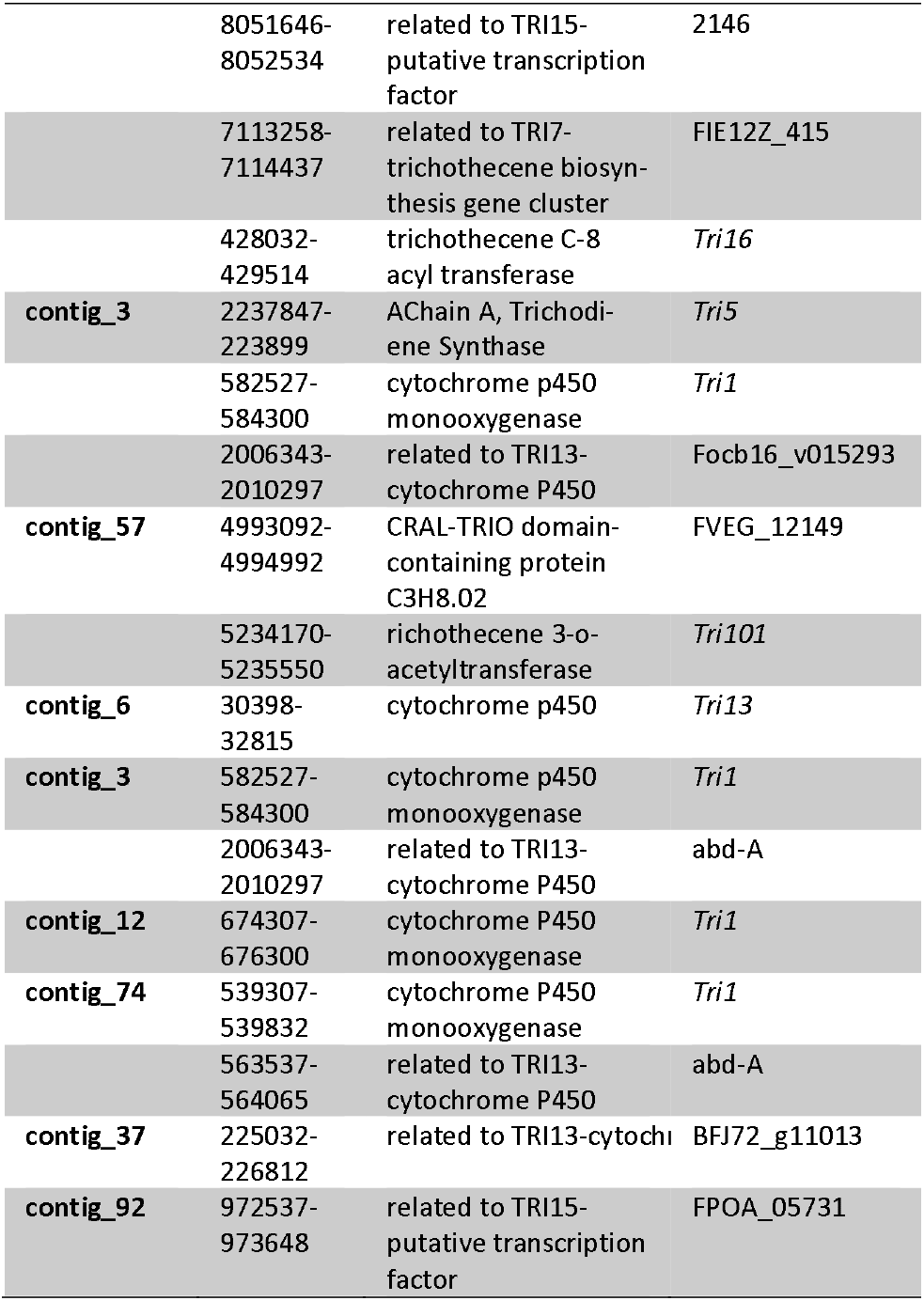
Gene position, for the TRI gene cluster.

### 3.1 *TRI* genes

Table 8 lists gene hits related to the *TRI* genes cluster. In the BLAST stage, 33 sequences were found. Most of them were gathered at the thirteenth contig of our assembly, while some were gathered at the fifty-third and fifty-seventh contigs amongst others.

### 3.2 HC-toxin related genes

Five proteins were found to be related to HC-toxin; all of them are listed in Table 9. Unlike the previous assembly, we identified two copies of HC-toxin synthetase located on the same contig and only 25 bases apart. Three proteins acted as an HC-toxin efflux carrier TOXA,

**Table 9:** Gene position, similarity gene and description of genes related with the keyword “hc-toxin” in assembly annotation.

| Contig | Position | Description | Gene |
| --- | --- | --- | --- |
| contig_11 | 265115-271850 | hc-toxin synthetase (non-ribosomal peptide synthetase) | KPA36315 |
|  | 271875-272769 | hc-toxin synthetase (non-ribosomal peptide synthetase) | KPA36315 |
| contig_13 | 989078-990512 | hc-toxin efflux carrier | Efflux pump roqT |
| contig_34 | 770120-771587 | hc-toxin efflux carrier | Efflux pump roqT |
| contig_93 | 1020288-1023864 | hc-toxin efflux carrier | Efflux pump roqT |

### 3.3 Related secondary metabolite genes

In addition, some contigs were linked to specific genes which are involved in global secondary metabolite biosynthesis. These include aldehyde reductase member 3 (Contig 13) and an efflux pump (Contig 2, 13, 57), ketose reductase (Contig 6, 13).

## 4 Discussions

With third-generation sequencing, the long reads and high depth generated using the PacBio^®^ SMRT sequencing led to a very high-quality assembly. The quality was not determined according to the contig length or N50 alone; BUSCO was another parameter used to examine the quality of the assembly. Compared to the publicly available assembly of Fl201059 which did not show good quality in the scaffold statistical data, but it had a high BUSCO compared to the *Saccharomyceta* dataset. Our assembly had both high-quality contigs and a high BUSCO rate, which means that more coding genes could be predicted by Augustus and GeneID. This provided hints to improve the accuracy of prediction. However, although the hints file could improve the accuracy, the core model in Augustus still came from *F. graminearum*, a related species. However, training Augustus produced a model based on the *F. langsethiae* sequence file. With the absence of an aligned BAM file or gene bank structure file from EnsembleFungi Fl201059, a FASTA file containing protein in the same sample was used to train the Augustus model. While the *ab initio* approach predicted genes with a model from a different species, it has less accuracy compared to the hints and training. Although the hints file improved the accuracy from *ab initio* to some extent, for some new species such as *F. langsethiae*, the accuracy from the hints file did not achieve the expectations of the analysis. Training Augustus with data that had 96% similarity with the assembly predicted more genes than *ab initio* and the hints file.

### Mycotoxin production pathways

The species in the *Fusarium* family are known to produce a wide range of secondary metabolites including type A and type B trichothecenes and zearalenone. Most significant strides have been made in relation to an understanding of the role of gene clusters involved in type B trichothecenes and fumonisins, which are produced predominantly by *F. graminearum*/*F. culmorum* and *F. verticillioides*/*F. proliferatum,* respectively (Munkvold, 2016). In addition, the former species also produces other mycotoxins, such as Zearalenone, Fusaric acids and Moniliformin especially in temperate cereals (Garcia-Cela et al., 2018). However, legislation on contamination of temperate cereals is predominantly focused on type B trichothecenes such as deoxynivalenol and zearalenone. Based on substitutions at C-8 and other positions around the core structure, trichothecenes have been classified into four groups (types A, B, C and D) (Munkvold, 2016), *Fusarium* species only produce type A and type B. Type A, which have an ester or hydroxyl or no oxygen substitution in the position of C-8, are usually produced by *F*. *armeniacum*, *F*. *langsethiae*, *F. poae*, *F*. *sambucinum*, *F*. *sporotrichioides*, and *F*. *venenatum*. Type B, which have a carbonyl group at C-8, are mainly produced by *F. graminearum* and *F. pseudograminearum* and *F. culmorum*. Type A compounds are generally more toxic than type B trichothecenes while the latter are usually produced in higher concentration. T-2 and HT-2 toxins are type A trichothecenes which mainly accumulate in oats and can cause immunological or haemotological defects in animals and potentially humans (CONTAM, 2011). Almost all proteins related to trichothecene biosynthesis are located on the 13^th^ contig of the assembly, with an additional copy of *TRI5* on contig 3 (See Fig. 1).

**Fig. 1.**
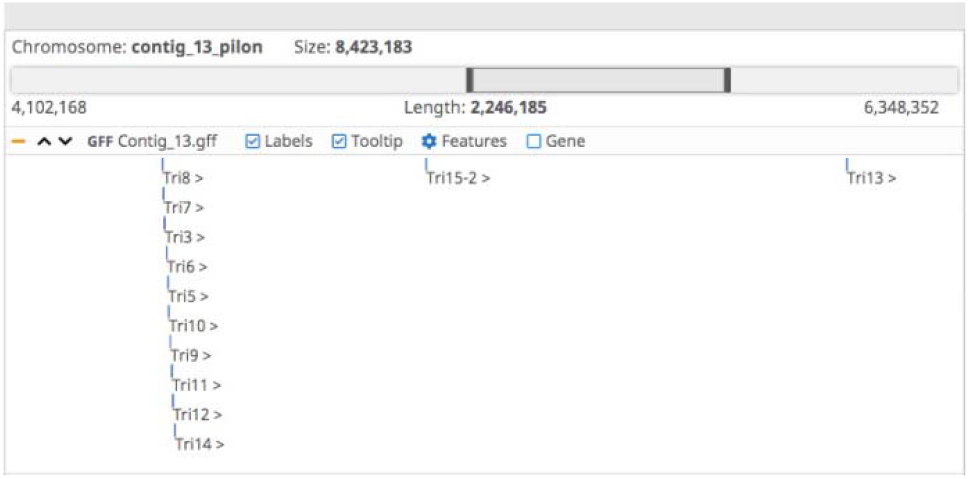
TRI-genes cluster identified on Contig 13 of the assembly.

Other proteins located in other contigs did not seem to have a core function in T-2 and HT-2 biosynthesis, and most of them encoded the transformation or production of TRI-proteins. HC-toxin synthetase was identified on contig 11 as two copies only 25 bps apart. Both copies were identical to the previous *F. langsethiae* assembly, but with 14 a.a. changes compared to the closest blast hit of the RGP60017 gene of *F. sporotrichioides* (See Fig. 2). It could be therefore inferred that contig 13 contains the main functional proteins regrouped in a cluster. However, one of the genes encoding a protein involved in HT-2 and T-2 mechanism called TRI1, were found in the contig 3, 12, 53 and 74.

**Fig. 2.**
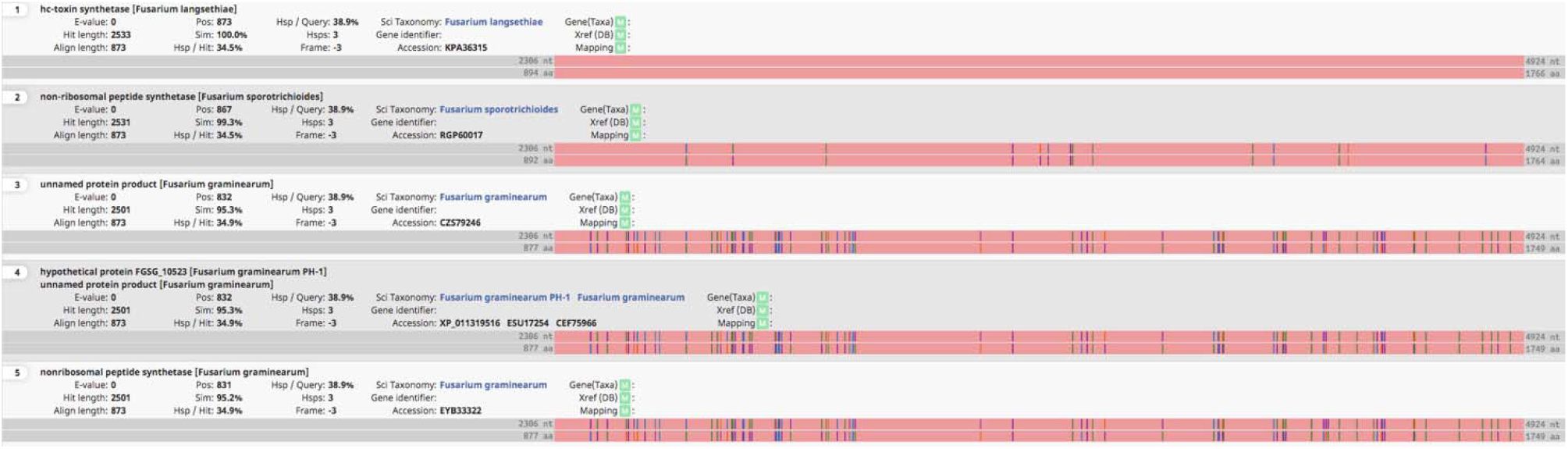
Blast hits of the hc-toxin gene highlighting 14 a.a.changes compared to the closest blast hit of F. sporotrichioides.

In 2011, McCormick described trichothecenes biosynthesis in the *Fusarium* species. Based on these findings, the biosynthesis process in *F. langsethiae* with different proteins could be inferred with the gene BLAST results and descriptions of proteins in *F. langsethiae* based on previous studies of other *Fusarium* species and the previous draft genome of *F. langsethiae* (Lysoe, 2016).

The first step consists in the cyclisation of farnesyl pyrophosphate, which is a primary metabolic intermediate (McCormick, 2011) and is mediated by a trichodiene synthase encoded in the gene *TRI5* in the 13^th^ contig as well as another hit identified on contig 3, suggesting its presence in two copies. *TRI5* is the core gene that mediates the biosynthesis of different trichothecenes (gi | 136010 | sp | P13513.1 | TRI5_FUSSP), including T-2 toxin. The ontology term of this gene (GO:0045482) indicates its molecular function as a trichodiene synthase. Tri5 is involved in the catalysis of the following reaction: 2-trans, 6-trans-farnesyl diphosphate = diphosphate + trichodiene. The *TRI5* gene was first characterised in a *F. sporotrichioides* strain that produced T-2 toxin (Hohn, 1986).

The trichodiene then goes through an oxygenation series catalysed by cytochrome P450 monooxygenase encoded by *TRI4* (McCormick, 2011). The *TRI4* gene (gi | 927758023 | gb | KPA41245.1) encodes a mono-oxygenase molecular function (GO:0004497) leading to the addition of four oxygens at C-2, C-3, C-11 and C-12, using C-13-epoxide to form the intermediate isotrichotriol (McCormick, 2006).

Subsequently, isotrichodermol (C-3-OH) is converted to isotrichodermin (C-3-OR) via an acetyltransferase encoded by *TRI101* (gi | 927756670 | gb | KPA40029.1) in the fifty-seventh contig (McCormick, 1999). The toxicity of *Fusarium* trichothecenes should be effectively reduced with this step, which serves as a mechanism for the fungal self-protection other trichothecene-producing organisms (Kimura, 1998). TRI101 (gene located in contig 57) acts as part of the transferase activity (GO:0016747) that transfers an acyl group, other than aminoacyl, from one compound to another. Then, a second hydroxyl group is added to C-15, which is controlled by TRI11 in the contig 13 (Alexander, 1998). TRI11 (gi | 927758018 | gb | KPA41240.1) works with a molecular function (GO:0016705)—an oxidation-reduction reaction in which hydrogen or electrons are transferred from each of two donors as well as an oxidation-reduction process (GO:0055114). After this TRI3 (gi | 927758024 | gb | KPA 41246.1) (GO:0043386) catalyses the acetylation of the 4-hydroxyl to form trichodermin and then, TRI13 protein (gi | 927758016 | gb | KPA41238.1) perform the same oxidation-reduction reaction as TRI11 in C-4 (Lee, 2002), followed by another acetylation process by TRI7 (gi | 927758025 | gb | KPA41247.1).

The next step of this process in *F. sporotrichioides* is the addition of a fourth hydroxyl group to C-8 by TRI1, followed by an addition of an isovaleryl moiety thanks to TRI16. Finally, the C-3 position loses the acetyl group via a TRI8-esterase step to produce T-2 toxin (McCormick, 2002).

In this study, the *TRI1* gene was found in the 3^rd^ and 53^rd^ contigs, and there was about 15% sequence dissimilarity with the *TRI1* sequence (gi | 927755786 | gb | KPA39264.1) in the database. Since *TRI16* was not found, it might have been either mis-matched and mis-labeled by BLAST or missing from the assembly. The gene *TRI8* was also found (gi | 927758026 | gb | KPA41248.1) and was identified as encoding a triglyceride lipase activity (GO:0004806) and is involved in a reaction in which triacylglycerol + H_2_O = diacylglycerol + a carboxylate.

Most of the genes linked to T-2 mycotoxin production were found in this assembly, but the *TRI9* gene only had sequences for which no literature could be found to describe their function. However, *TRI9* has been found not only in *F. langsethiae* but also other *Fusarium* species, such as *F. sporotrichioides* and *F. graminearum*. It acts upon the integral component of the membrane (GO:0016021) and might be linked with the T-2 mycotoxin transport mechanism.

With regard to the genes identified in relation with the HC-toxin, not enough information was found to support the production pathways in *F. langsethiae*. However, some genes encoded global regulators such as the efflux pump and carrier of HC-toxin produced by *Cochliobolus* species might have an evolving relationship with *F. langsethiae*.

## Availability

This Whole Genome Shotgun project has been deposited at GenBank under the accession JAFFKB000000000 under BioProject PRJNA701381. The version described in this paper is version JAFFKB010000000.

